# Identification of a new population of myonuclei during skeletal muscle hypertrophy

**DOI:** 10.64898/2026.05.05.723044

**Authors:** Léa Delivry, Stéphanie Backer, Maxime Di Gallo, Alix Silvert, Mathieu Dos Santos, Florian Britto, Pascal Maire, Athanassia Sotiropoulos

**Affiliations:** Université de Paris Cité, Institut Cochin, Inserm, CNRS, Paris, France; Université de Paris Cité, Paris, France; University of Texas Southwestern Medical Center, Dallas, TX, USA

**Keywords:** Muscle hypertrophy, plantaris overload, snRNA-seq, RNAscope, RhoA, Postn, Gsn

## Abstract

**Background:** Skeletal muscle represents around 40% of total human body weight and exhibits remarkable plasticity. It can hypertrophy, atrophy, or regenerate in response to changes in activity, nutrient availability, or injury. The main component of striated muscle, the myofiber, is a post-mitotic, multinucleated cell that contains the muscle’s contractile unit, the sarcomere. The myonuclei within these fibers are specialized and differ in terms of gene expression and localization. Adult muscles also contain various other cell types, including adult muscle stem cells (MuSCs), macrophages, fibro-adipogenic progenitors (FAPs), and endothelial cells. MuSCs are central to muscle plasticity, and are capable of activation, proliferation, differentiation, and fusion to form new myofibers during regeneration, or to fuse with existing myofibers during hypertrophy. Muscle hypertrophy and myofiber’s enlargement involve increased protein synthesis and reduced protein degradation, as well as myonuclear accretion following satellite cell activation. Multiple signaling pathways, such as the mTOR pathway and the RhoA/SRF mechanotransduction pathway, are involved in these processes.

**Methods:** We performed single-nucleus RNA sequencing (snRNA-seq) on *plantaris* muscles of adult mice, comparing samples 7 days after hypertrophy induction (overload, 7OV) to non-hypertrophied controls (Ctl). RNAscope experiments on isolated myofibers identified the heterogeneity of myonuclei along the myofiber.

**Results:** SnRNA-seq analysis revealed a previously unknown population of myonuclei (UM). UM-Ctl, which is present only in the Ctl condition, and UM-7OV, only in the 7OV condition. These myonuclei are localised at the tips of myofibres. Furthermore, we determined that UM-7OV are not newly fused myonuclei from activated satellite cells. Trajectory analyses suggest that UM-Ctl transition into UM-7OV during hypertrophy, returning to a near-basal homeostatic state after 21 days of overload (21OV). Gene expression analysis showed that UM-Ctl and UM-7OV have distinct gene expression profiles compared to other myonuclei and respond differently to hypertrophy.

**Conclusion:** Our findings suggest the existence of a specific population of myonuclei with unique localization and gene expression profiles, which play distinct roles at baseline and during hypertrophy. These results highlight the differential properties of myonuclei in the myofiber and their potential specific functions in muscle homeostasis and adaptation.

## Background

Skeletal muscle, one of the most abundant tissues in the vertebrate body, possesses a high degree of plasticity and can adapt its size, energy metabolism and fibre phenotype to workload. Adult skeletal muscle tissue is composed of several cell types, including vessel-associated cells, myofibroblasts and the resident muscle stem cells (also known as muscle satellite cells, MuSCs). However, the most important cell type of skeletal muscle in terms of tissue locomotion function and abundance is the multinucleated myofiber. Physiological demands such as resistance exercise or functional overload (OV) lead to an increase in adult muscle mass due to a hypertrophic growth of myofibers. The two main mechanisms involved in muscle growth are: i) the regulation of proteostasis towards enhanced protein synthesis including ribosome biogenesis at the expense of protein degradation to supply new contractile filaments to growing myofibers, and ii) the fusion of new nuclei provided by MuSCs, leading to the accretion of new myonuclei which is required for muscle hypertrophy [1] and may increase the transcriptional and translational capacity of the syncytium [2,3]. Indeed, in response to increased workload, MuSCs exit the quiescent state, proliferate, differentiate and subsequently fuse with growing myofibers [4]. Strikingly, MuSCs can also fuse with growing myofibers without a prior proliferation phase, as recently reported [5]. Nevertheless, the activation of satellite cells is necessary for the hypertrophy process [6].

The myofiber syncytium, which extends between tendons, contains hundreds of myonuclei that control their own myonuclear domain [7,8]. Despite the central role of myofibers in muscle biology, the specific functions of individual myonuclei within the myofiber have been largely overlooked. Recent single-nucleus RNA sequencing (snRNA-seq) experiments, which allow the capture of the transcriptional signatures of myonuclei, have highlighted the transcriptional heterogeneity between myonuclei and revealed the molecular signatures of subpopulations of myonuclei within a myofiber, including those of the neuromuscular junction (NMJ) and myotendinous junction (MTJ) domains [9–11] This myonuclear heterogeneity is associated with 3D chromatin heterogeneity among myonuclei has been revealed by snATAC-seq experiments [9,12,13]. However, how myonuclei heterogeneity may contribute to muscle plasticity during hypertrophic conditions remains to be addressed.

Adult skeletal muscle hypertrophy is controlled by several signaling pathways within myofibers, the most prominent of which is the mTOR pathway [2,14]. In response to increased load and to growth factors (IGF1, MGF), PI3K/Akt/mTOR signaling increases protein synthesis while inhibiting FOXO-dependent transcription and protein breakdown [15]. In addition, several mechanotransduction pathways are implicated in muscle growth. The small GTPase RhoA is capable of translating the physical forces into biochemical signaling and activating specific transcription factors including Serum Response Factor (SRF) [16–18]. Importantly, RhoA and SRF, within myofiber, are functionally involved in muscle hypertrophic growth, even though they are not necessarily directly linked [18,19] [20]. Yes-Associated Protein (YAP), another key player in mechanotransduction, has also been shown to play a role in hypertrophic growth [20]. The main goal of this study is to unravel the transcriptional signature of myonuclei within myofiber syncytium and to investigate the heterogeneity of myonuclei in response to overload in mice. We used as a paradigm the compensatory hypertrophy of the *plantaris* muscle following incapacitation of the synergistic muscles (*soleus* and *gastrocnemius*) (OV). To this end, we analyzed the transcriptional identity of each individual nucleus within the *plantaris* at steady state (Ctl) or after 7 days of OV using an snRNA-seq approach. We identified two categories/clusters of unknown myonuclei (UM-Ctl and UM-7OV) that are specific to Ctl and 7OV conditions, respectively. Having identified the specific markers of UM-Ctl (*gelsolin, Gsn*) and UM-7OV (*periostin, Postn*) myonuclei, we showed their specific localisation at the tips of myofibers. Furthermore, we highlighted a high degree of transcriptomic similarity between these two clusters, suggesting that UM-Ctl myonuclei become UM-7OV myonuclei in response to hypertrophy. In addition, we provide several lines of evidence that argue against UM-7OV myonuclei representing newly fused myonuclei provided by MuSCs. Finally, Gene Ontology analysis suggested that UM myonuclei respond differently to OV compared to the other myonuclei, highlighting the striking heterogeneity in the transcriptional response of myonuclei to hypertrophy within the myofiber.

## Methods

### Overload-induced hypertrophy

Overload-induced hypertrophy (OV) of *plantaris* muscles of 8-10 week-old female WT(C57Bl6N) mice, 12-16 week-old male and female *ROSA^mTmG^;PAX7^CreERT2/+^* (mTmG) mice [21,22] and 8-10 week-old female *RhoA^flox/flox^;HSA-CreERT2* mice [18] was induced through sectioning the tendon of soleus and gastrocnemius muscles. This procedure was achieved in both legs. Plantaris muscles were dissected and subsequently processed for histological analyses. *RhoA^flox/flox^;HSA-CreERT2* mice were infected intraperitoneally with 100µL (concentration 20µg/µL) of Tamoxifen during 5 days (MP Biomedicals, 156738).

All animal experiments were conducted in accordance with the European guidelines for the care and use of laboratory animals and were approved by the institutional ethic committee and FrenchMinistry of Research (number A751402).

### FACS and Single nucleus RNAseq sequencing

For each condition, 12 *plantaris* were used. *Plantaris* muscles were prepared according to the STAR Protocol « Extraction and sequencing of single nuclei from murine skeletal muscles », 2021. Nuclei were then FACS sorted to exclude debris with a BD FACSAria III and the BD FACSDIVA software. (https://support.10xgenomics.com/single-cell-gene-expression/software/pipelines/latest/advanced/references). We used a reference genome built against mouse mm10, Sequence: GRCm38 Ensembl 93 and Cellranger v6.0.0 and v7.1.0.

### Single nuclei RNAseq processing

For the bioinformatics analyzes we used R (v4.2.0) and Seurat (v4.2.0). We obtained between 3500 and 4000 nuclei per condition. Scripts are available on our team Github: https://github.com/mairesotiropoulos/Single-nucleus-RNA-seq/Hypertrophy

### FISH with amplification (RNAscope) on single myofibers

RNAscope® Multiplex Fluorescent Assay V2 (ref 323100) was used to visualize *Postn* (Mm-Postn-O1-C2,1123651-C2), *Gsn* (Mm-Gsn-O1-C3, 124047-C3), *Ttn* (Mm-Ttn-intron1, 824501) pre-mRNAs and *Postn* mRNAs (Mm-Postn-C2, 418581-C2). *Plantaris* muscles were dissected from the leg with their tendons. Muscles were fixed with PFA 4% (Electron Microscopy Sciences,15735-90) during 30min and washed 3×10min with PBS 1X at 4°C. The fixed muscles were then dissected in PBS 1X with two needles to obtain single fibers. Single fibers were then fixed on Superfrost plus slides (Thermo Fischer) coated with Cell-Tak (Corning) by dehydration at 55°C during 5 min. Myofiber were treated according to the manufacturer’s protocol: ethanol dehydration and dry, hydrogen peroxyde, protease IV, hybridation with probes for 2h and revelation treatment. Slides were were mounted under a glass coverslip with Mowiol 4-88 (Calbiochem). Myofibers were imaged with a Leica DMI6000 confocal microscope composed by an Okogawa CSU-X1M1 spinning disk and a CoolSnap HQ2 Photometrics camera. Images were analyzed with Fiji Cell counter program.

### Trajectory PAGA pipeline

To obtain a trajectory of our data we used the PAGA pipeline on git-hub: https://github.com/theislab/paga.

### GO analysis

We used the software Ingenuity (IPA): https://digitalinsights.qiagen.com/products-overview/discovery-insights-portfolio/analysis-%20and-visualization/qiagen-ipa/.

## Image acquisition

All the fluorescence image were acquired with a Spinning Disk confocal microscope (Yokogawa X1) with 10X (Immunofluorescence) or 20X (RNAscope) objective (HCX PL APO, NA 1.47).

## Quantification and statistical analysis

All the quantitative data sets were analyzed using a paired parametric t-test with Welsh correction (Figures 2D, 2G, 2K, 5C, 5E, 5F, 5H), and an unpaired parametric t-test with Welsh correction (Figure 2F, 2J) using GraphPad Prism 8.4.0 software. Statistical significance was set at a p value<0.05.

## Results

### Unknown myonuclei (UM) identified in Ctl and 7OV *plantaris* muscles

In order to investigate the transcriptomic signature of myonuclei during hypertrophy, we performed snRNA-seq on *plantaris* muscles at basal state (Control, Ctl) and 7 days after OV-induced hypertrophy (7OV). Two independent experiments were conducted using two mouse models: C57Bl/6N mice (WT) and *ROSA^mTmG^;PAX7^CreERT2/+^* (mTmG) mice. Following *plantaris* dissection and nuclei FACS sorting using DAPI, we carried out snRNA-seq experiment using 10X technology on the collected nuclei. Nuclei from Ctl and 7OV muscles were sequenced and analysed. After pooling these two experiments, the Ctl and 7OV data were then integrated and a Uniform Manifold Approximation and Projection (UMAP) was obtained with 3972 nuclei for the Ctl condition and 2997 nuclei for the 7OV condition (Figure 1A). Clusters corresponding to different cell types were identified based on the differential expression of canonical markers such as *Pax7* (MuSCs, adult muscle stem cells), *Ptprc* (macrophages), *Pdgfra* (FAPs, fibroblastic adipocyte progenitors), *Col11a1* (tenocytes), Pecam1 (endothelial cells) and *Myh11* (smooth muscle cells), as previously described [9]. Myonuclei expressing *Titin* (*Ttn*) were separated into different clusters: *Myh1* (type IIx heavy chain Myosin), *Myh4* (type IIb heavy chain Myosin) and *Myh2* (type IIa heavy chain Myosin) clusters identified by their Myosin expression, the MTJ (myotendinous junction) cluster identified by *Col22a1* expression and an unknown cluster designated as UM (Unknown Myonuclei) (Figure 1A - Figure S1-S2). This unknown myonuclear cluster (UM) expressed *Ttn*, *Myh1* and *Myh4*, as did the other myonuclei. However, it exhibited reduced expression of *Ttn* compared to the body myonuclei (*Myh1*, *Myh2* and *Myh4* clusters). The UM cluster also overexpressed *Meg3* and *Neat1*. Its distinct transcriptional signature and isolation from the other myonuclei clusters suggested that this cluster contained myonuclei distinct from the rest of the body myonuclei. Importantly, similar clusters were obtained when the datasets from the two independent experiments were analyzed separately, demonstrating the reproducibility of UM-Ctl and UM-7OV cluster emergence (Figure S3).

**Fig. 1.**
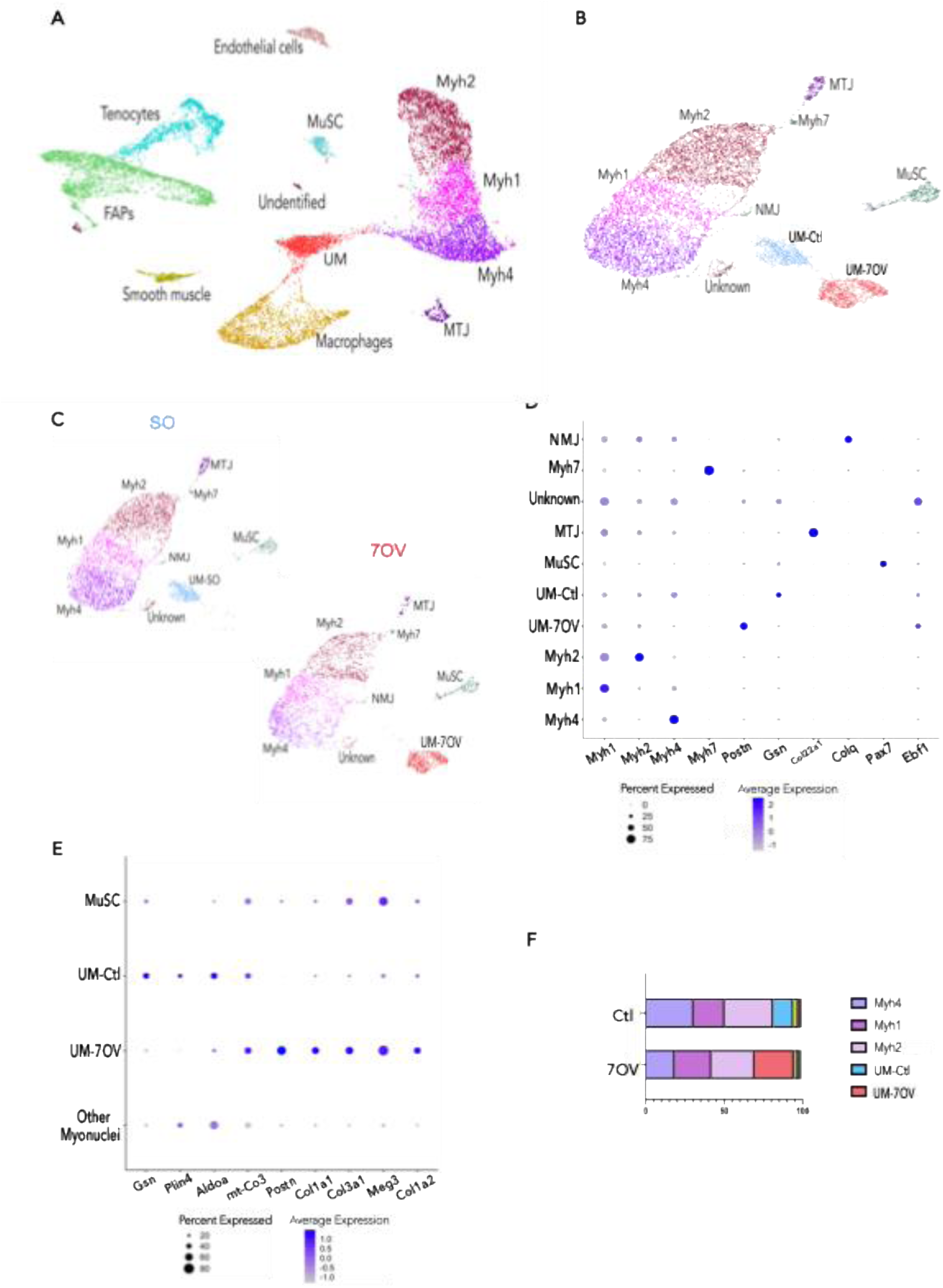
Unknown populations of myonuclei are present in Ctl and 7OV plantaris muscle. **A** Uniform Manifold Approximation and Projection (UMAP) visualization represents all the nuclei present in Ctl and 7OV plantaris muscle of WT and mTmG mice. **B** UMAP with the reclustering of myonuclei only present in Ctl and 7OV plantaris muscle of WT and mTmG mice. **C** Split UMAP with only Ctl myonuclei or 7OV myonuclei in plantaris muscle of WT and mTmG mice. **D** Dotplot depicting biomarkers of each myonuclei cluster. **E** Dotplot depicting biomarkers of MuSC, UM-Ctl, UM-7OV and other myonuclei clusters. **F** Graph representing the proportion of myonuclei subtypes in Ctl and 7OV conditions.

**Fig. 2.**
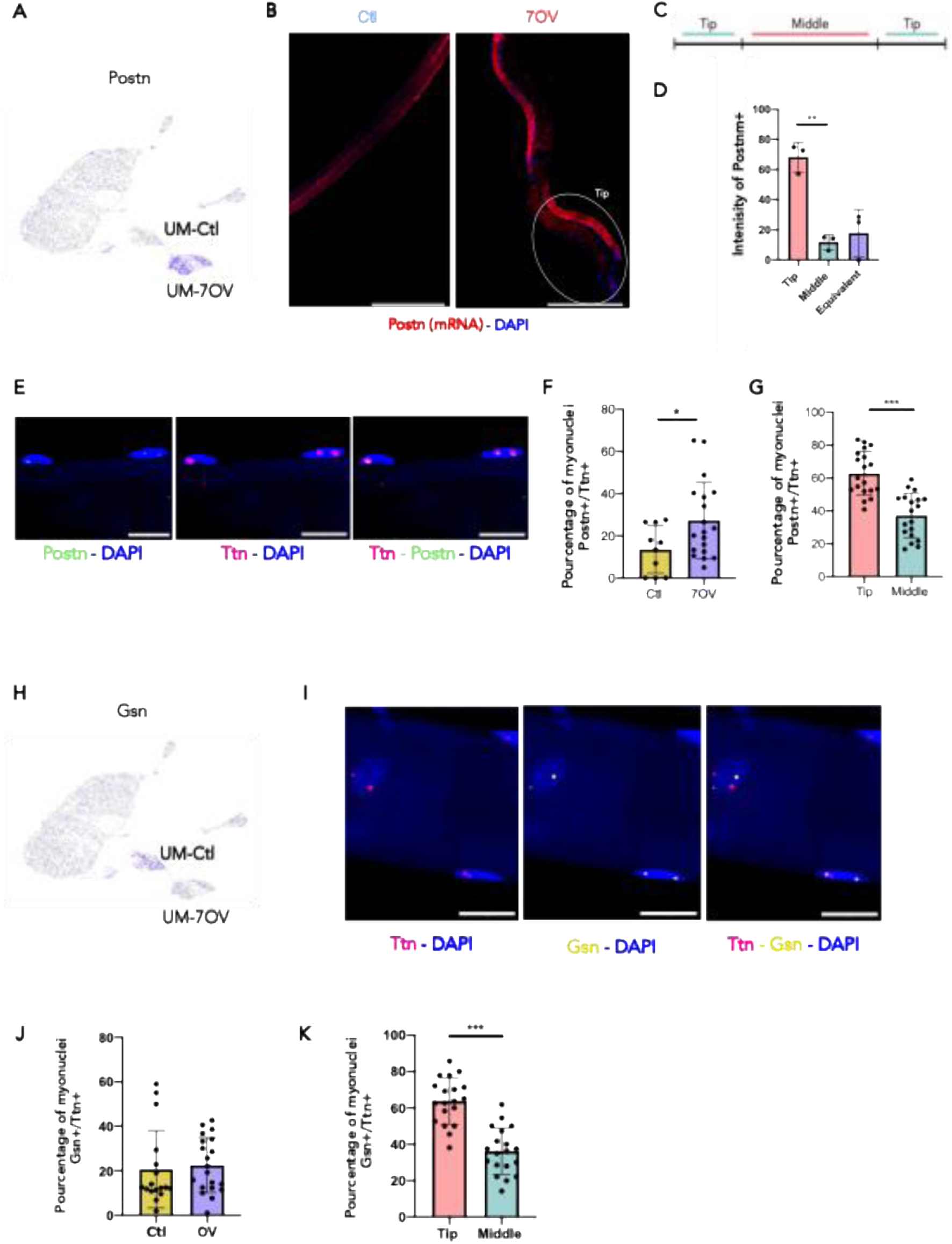
Localization of UM-Ctl and UM-7OV myonuclei at the tips of the myofibers. **A** Dimplot with *Postn* expression in UM-7OV cluster from Fig. 1B. **B** Plantaris myofiber of Ctl and 7OV mice hybridized with *Postn* mRNA (red) and DAPI (blue). Scale = 50µm. **C** Representation of myofiber cut levels for counting. **D** Localization of the intensity of *Postn* mRNA in 7OV myofibers (n=30 fibers, N=4 mice). **E** Plantaris myofiber hybridized with *Postn* pre-mRNA (green), *Ttn* (red) and DAPI (blue) in 7OV mice. Scale = 20µm. **F** Number of *Postn+/Ttn+* myonuclei per fiber of Ctl and 7OV mice (N=3 mice Ctl and N=3 mice 7OV). **G** Localization of myonuclei Postn+/Ttn+ in 7OV myofibers (n=19 fibers, N=4 mice). **H** Dimplot with *Gsn* expression in UM-Ctl cluster from Fig. 1B. **I** Zoom of a plantaris myofiber hybridized with *Gsn* pre-mRNA (yellow), *Ttn* (red) and DAPI (blue) in Ctl mice. Scale = 20µm. **J** Percentage of *Gsn+/Ttn+* myonuclei per fiber in Ctl and 7OV mice (N=3 mice Ctl and N=4 mice 7OV). **K** Localization of *Gsn+/Ttn+* myonuclei in Ctl myofibers (n=17 fibers, N=4 mice).

To investigate more specifically the effect of hypertrophy on the transcriptional landscape of myonuclei, myonuclei and MuSC nuclei were reclustered. The four myonuclei sub-clusters Myh1, Myh2, Myh4 and MTJ and the MuSC cluster remained unchanged compared to the clustering involving all muscle nuclei (Figure 1A-B). However, new clusters were detected: the cluster expressing *Myh7* (type I Myosin heavy chain), the neuro-muscular junction (NMJ) cluster expressing *Colq* and an unknown cluster expressing *Ebf1* (Figure 1B-1D - Figure S4-S5). In addition, the UM cluster was again clustered separately from the other myonuclei (Myh1, Myh2, Myh7, Myh4, MTJ and NMJ clusters), but was now divided into two sub-clusters: the UM-Ctl and the UM-7OV. UM-Ctl cluster is characterised by some biomarkers like *Gsn, Plin4 and Aldoa* and expressed more *Myh4* than other *Myosin heavy chain* genes (Figure 1B-1D-1E - Figure S4-S5). And the UM-7OV cluster is characterised by biomarkers like *Postn, Meg3, Col3a1, Col1a1* and expressed more *Myh1* than other *Myosins* (Figure 1B-1D-1E - Figure S4-S5). Strikingly, UM-Ctl cluster was only present in Ctl samples and UM-7OV cluster was only present in OV conditions (Figure 1C). Analysis of the percentage of nuclei in each cluster confirmed the strong reduction of the UM-Ctl population (from 13% in Ctl condition to 0.7% in OV) and the appearance of UM-7OV after OV (from 0.2% in Ctl to almost 25% in OV) (Figure 1F).

Notably, Ctl myonuclei expressed more *Myh4* (30.4% of nuclei) than 7OV myonuclei (18.5%), but less *Myh1* (19.9% vs 23.5%) (Figure 1D-1F), confirming the switch from fast glycolytic *Myosins* to slower oxidative Myosins during skeletal muscle hypertrophy (Figure 1D-1F) [23,24]. In addition, the *plantaris* muscle is a fast muscle with many hybrid fibres expressing two types of *Myosin heavy chain* (*Myh*) genes and proteins within the syncytium [25]. Myonuclei expressing two *Myh* with an expression per cell >3 were quantified to determine the number of hybrid myonuclei. During hypertrophy, the number of myonuclei expressing two *Myh* increased in our data: from an average of 5% of hybrid myonuclei co-expressing *Myh1* and *Myh2* in the Ctl condition to an average of 10% in the 7OV condition. This result is again consistent with the shift of *Myh* during hypertrophy. The number of hybrid myonuclei expressing *Myh4* and *Myh1* remained virtually unchanged (4% for Ctl myonuclei vs. 2% for 7OV myonuclei).

We compared the biomarkers of the UM-Ctl and UM-7OV with those reported in the literature for various myonuclei subpopulations. We hypothesized that the UM cluster might correspond to the MTJ-B cluster expressing *Col6a1, Col6a3 and Ebf1* described by Birchmeier’s team [10] but they do not correspond to them (Figure 1E). Furthermore, by comparing with the unknown myonuclei from the Dos Santos study (expressing *Sgms2, Myh9, Dysf, Nrap, FlnC* and *Runx1*, which are associated with membrane repair and myofibrillogenesis processes) and the MTJ-Bs from the Kim study to UM-Ctl, we concluded that UM-Ctl did not correspond to either of these two types of myonuclei [9–11].

Taken together, these data reveal two distinct subpopulations of myonuclei (UM-Ctl and UM-7OV), which are specific for two different functional states of muscle: at rest and during an increased workload.

### UM-Ctl and UM-7OV are localised in the tips of myofibers

The next step was to further characterise these two unknown myonuclei subpopulations by investigating the location of UM-Ctl and UM-7OV within a myofiber. To this end, we performed RNA-FISH experiments (RNAscope) on isolated single fibres using a pre-mRNA probe against *Gsn* and mRNA and pre-mRNA probes against *Postn* to visualise UM-Ctl and UM-7OV, respectively (Figure 2A-2H).

In an attempt to localise UM-7OV, a probe against *Postn* mRNA was used to determine if and where *Postn* messengers accumulate in the myofiber syncytium during hypertrophy. As shown in Figure 2B, very little *Postn* mRNA expression was detected in Ctl myofibers (Figure 2B, left panel), whereas high levels of *Postn* mRNA were observed 7 days after OV (Figure 2B, right panel), which is consistent with the snRNA-seq data (Figure 2A). Notably, all single myofibers expressed *Postn* after OV, suggesting that UM-7OV are present in all myofibers and are not restricted to a few specific myofibers expressing *Postn*. To determine the location of the mRNAs of interest, the myofibers were virtually separated into middle and tip regions (Figure 2C). Interestingly, after 7OV, more than 65% of the myofibers showed a preferential accumulation of *Postn* mRNAs in their tips, 15% in the middle part and 20% had a uniform *Postn* distribution (Figure 2D). To specifically label UM-7OV myonuclei, we repeated RNA-FISH experiments with *Postn* and *Ttn* pre-mRNA probes. *Ttn* staining was used to distinguish myonuclei from nuclei of other cell types (Figure 2E). First, we quantified the proportion of *Postn+/Ttn+* myonuclei relative to the total number of *Ttn+* myonuclei before (Ctl) and after seven days of OV (7OV). As with *Postn* mRNA labelling and snRNA-seq data, an increase in the number of *Postn+/Ttn+* myonuclei was observed in the OV condition compared to the Ctl condition (Figure 2F). Furthermore, consistent with *Postn* mRNA RNAScope data, *Postn+/Ttn+* myonuclei were preferentially localised to the tips of 7OV myofibres (Figure 2G).

A similar RNAscope approach was performed to quantify and localise UM-Ctl using a pre-mRNA probe against *Gsn* as a marker (Figure 2H-2I). The proportion of *Gsn+/Ttn+* myonuclei is also the same in Ctl and OV conditions (Figure 2J). Similar to *Postn+/Ttn+* UM-7OV in OV conditions, *Gsn+/Ttn+* UM-Ctl in Ctl conditions were also preferentially localised to the tips of Ctl myofibers (Figure 2K). Altogether, these results suggest that within the myofiber syncytium, there is a subpopulation of myonuclei (UM) that are preferentially located at the tips in both the basal state (UM-Ctl) and after OV (UM-7OV).

### RhoA is essential for the maintenance of UM-Ctl and UM-7OV in myofibers

Interestingly, UM-Ctl and UM-7OV are preferentially located at the ends of the myofibers close to the tendons, a region of the myofibers exposed to high tension/force [26]. We therefore hypothesised that these myonuclei might be involved in mechanical sensing and possibly in mechanotransduction signaling. The RhoA pathway has been identified as an important player in mechanotransduction [16] and we have previously shown that RhoA in myofibers is required for overload-induced hypertrophy [27]. We therefore wondered whether RhoA signalling might affect the UM-Ctl and UM-7OV populations. To this end, we performed similar snRNA-seq experiments on Ctl and 7OV *plantaris* muscles from *RhoA^flox/flox^;HSA-CreERT2* (hereafter referred to as mutant - Mut) mice, in which *RhoA* was deleted in myofibers after tamoxifen injection (Figure 3A). We integrated this new data with that used in Figure 1. We then performed a reclustering and obtained a UMAP that identified the same clusters as previously described (Figure 3B). Surprisingly, when we looked at the expression of UM-Ctl and UM-7OV biomarkers, we saw a decrease in expression and in the number of myonuclei expressing these genes in the myonuclei of MutCtl and Mut7OV mice compared to mTmG and WT (Ctl and 7OV) mice (Figure 3C). The analysis of MutCtl and Mut7OV separately revealed that both UM-Ctl and UM-7OV subpopulations were almost absent in RhoA-mutant muscles. UM-Ctl represented only 0,7% of myonuclei in the MutCtl condition, while UM-7OV represented only 0,9% of myonuclei in the Mut7OV condition. Interestingly, UM-Ctl were found in the Mut7OV condition, representing 3% of their myonuclei (Figure 3D). Furthermore, the expression of *Gsn* in the body myonuclei (Myh1, Myh2 and Myh4 clusters) was similar in MutCtl and Ctl conditions (Figure 3E). *Postn* expression was also similar in body myonuclei in the 7OV and Mut7OV conditions (Figure 3F). These data suggest that UM-Ctl and UM-7OV, which overexpress *Gsn* and *Postn* respectively, are absent in mutants and are not dispersed in the other clusters of myonuclei.

**Fig. 3.**
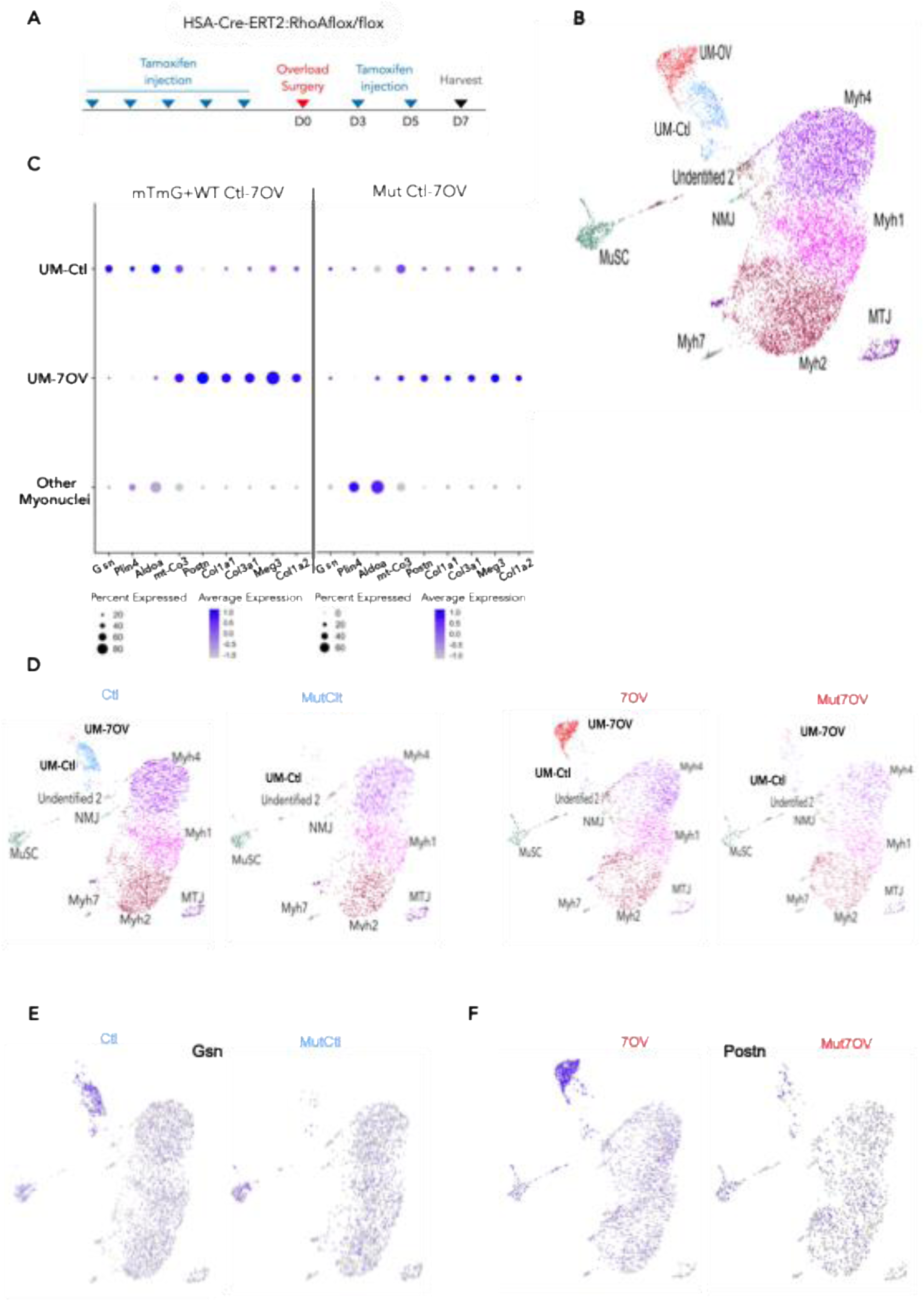
UM-Ctl and UM-7OV myonuclei are absent in *RhoA* mutant muscles. **A** *RhoA^flox/flox^ ;HSA-CreERT2* mice underwent surgery at D0, received daily tamoxifen injections for 5 consecutive days and were harvested at D7. **B** UMAP with the reclustering of myonuclei only present in Ctl, 7OV, MutClt and MutOV plantaris muscle. **C** Dotplot of markers from UM-Ctl, UM-7OV and other myonuclei clusters under Ctl and 7OV conditions and under MutCtl and Mut7OV conditions. **D** Split UMAP with only Ctl ,7OV, MutCtl and Mut7OV nuclei in plantaris muscle. **E** Dimplot with the expression of Gsn in Ctl and MutCtl plantaris muscle. **F** Dimplot with the expression of *Postn* in 7OV and Mut7OV plantaris muscle.

Therefore, RhoA may play a role in the maintenance of UM-Ctl and UM-7OV in the homeostatic muscle and in the 7OV condition.

### UM evolves between different states during muscle hypertrophy

We next wondered whether UM-Ctl and UM-7OV were related and whether they might represent adaptations of the same subpopulation of myonuclei in response to different physiological states, namely basal and increased workload. To test this hypothesis, we performed trajectory analyses on the myonuclei and MuSC from Figure 1B using the PAGA pipeline (Figure 4A). Interestingly, the UM-Ctl and UM-7OV clusters were directly connected, whereas the MuSC cluster was separated. Furthermore, UM-7OV was only connected to the UM-Ctl cluster, highlighting their transcriptional proximity even when the two clusters are not present under the same physiological conditions (Ctl vs 7OV).

**Fig. 4.**
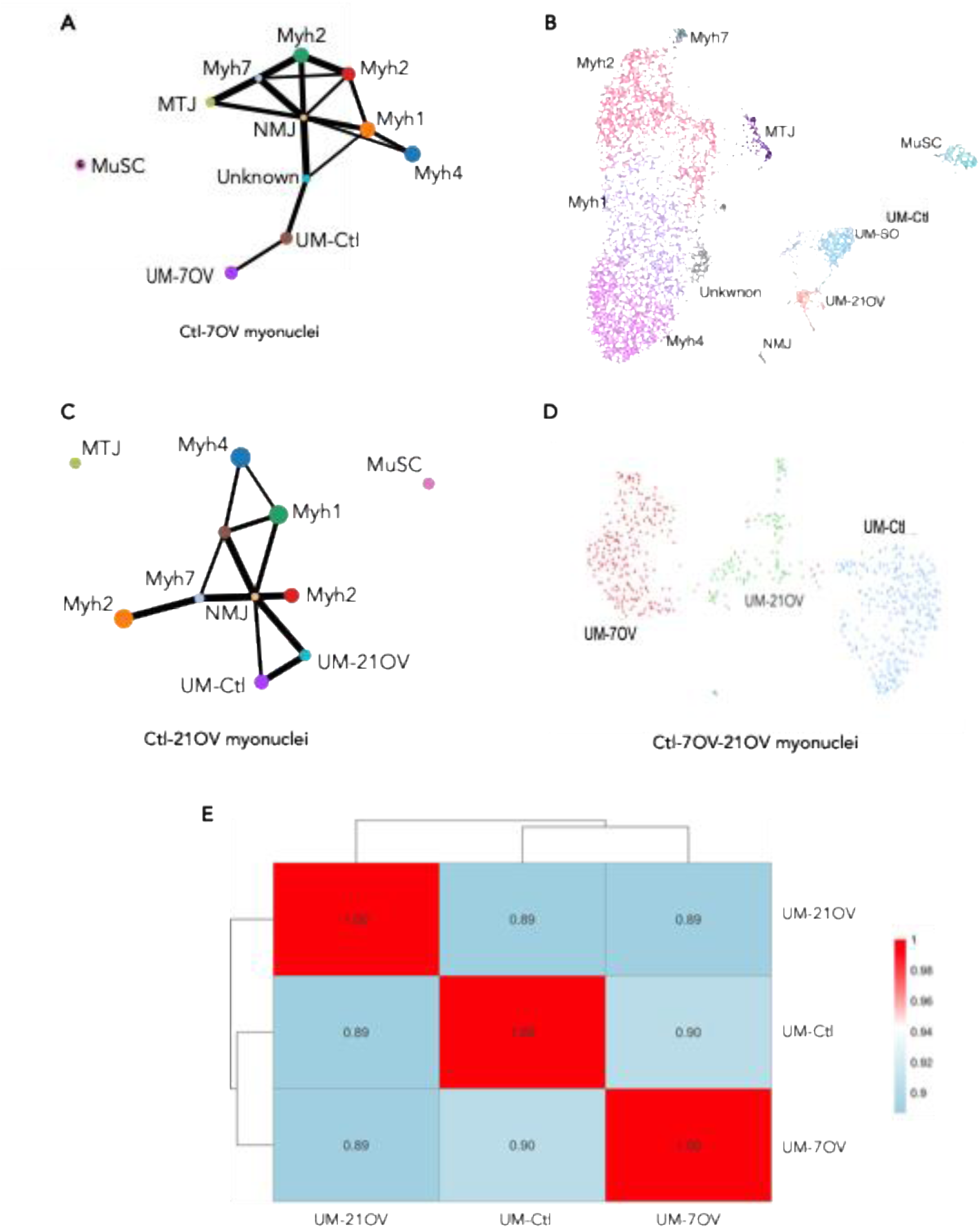
UM transitions between distinct states during skeletal muscle hypertrophy. **A** Trajectory of Ctl and 7OV myonuclei and MuSC during plantaris hypertrophy. **B** UMAP with only myonuclei of Ctl and 21OV muscle. **C** Trajectory of Ctl and 21OV myonuclei and MuSC. **D** UMAP with only Ctl, 7OV and 21OV UM in mTmG mice. **E** Transcriptional similarity between UM-Ctl, UM-7OV and UM-21OV.

To gain insight into the temporal evolution and fate of the UM-7OV population, we generated snRNA-seq data from mTmG mice after 21 days of OV (21OV) and integrated them with data from mTmG mice in the Ctl condition (Figure 4B). UMAP of the merged Ctl and 21OV mTmG myonuclei data revealed two UM cluster composed of a small amount of UM-21OV (only 1.8% of 21OV myonuclei) compared to the UM-Ctl cluster composed of Ctl myonuclei (18.8% of Ctl myonuclei). Interestingly, UM-21OV cluster did not express markers similar to those of the UM-7OV cluster, in particular *Postn*. Furthermore, PAGA trajectory analysis showed that the two clusters (UM-21OV and UM-Ctl) are connected, indicating that they are potentially transcriptionally close (Figure 4C).

We then compared the expression profiles of UM-Ctl, UM-7OV and UM-21OV. We took the Ctl, 7OV and 21OV myonuclei data from the mTmG mice and merged them. We then reclustered only the single UM cluster obtained by merging these three datasets. This reclustering resulted in three clusters corresponding to UM-Ctl, UM-7OV and UM-21OV (Figure 4D). We then performed an analysis of the transcriptional similarity between those three clusters (Figure 4E), showing the correlation between the average expression profiles of the selected clusters. A correlation below 0.50 suggests transcriptionally distinct cell types, while a correlation above 0.70 indicates similar cell types, possibly in different states or closely related subtypes. A high correlation (r = 0.90 and r = 0,89, r > 0,50) was observed between the average expression profiles of UM-Ctl, UM-7OV and UM-21OV clusters, suggesting that they may represent the same cell type but in different states (Figure 4E).

Taken together, these analyses suggest that UM-Ctl myonuclei, which are present at rest, respond transcriptionally to the hypertrophic process (7 days OV) and become UM-7OV by changing their gene expression. Then, at a later time point after OV (21 days), the muscle reaches a new homeostatic state that differs from the resting state (Ctl). The UM-21OV cluster represents an intermediate state from UM-7OV to UM-Ctl.

### Newly fused myonuclei are not included in UM-7OV sub-population

During hypertrophy, the number of myonuclei in a myofiber increases due to the activation of MuSCs and their subsequent fusion with the growing myofibers. We therefore wondered whether the UM-7OV cluster could correspond to newly fused myonuclei.

In a recent study, Millay’s team examined the bulk transcriptomes of sorted newly fused and pre-existing myonuclei after overload-induced hypertrophy (7 days) [28] and showed that expression of *H19, Myh3, Myh8, Tnnt2, Tnnt1* and *Postn* was enriched in newly fused myonuclei. This study was complemented by snRNA-seq experiments on FACS sorted myonuclei after overload-induced hypertrophy (7 days), which identified three myonuclear subpopulations modulated by OV (Unclassified A, B and *Atf3+* clusters) (Figure 5A, [28]). The authors attributed two of these clusters, Unclassified A and B, to newly fused myonuclei [28]. Unclassified A and B clusters both expressed *H19, Igf2, Runx1, Myh3* and *Myh8*, while *Tnnt2* and *Dclk1* expression was more specific for the Unclassified cluster B. We therefore wondered whether UM-7OV might represent an overlapping myonuclei population with these newly fused myonuclei represented by the two Unclassified clusters A and B. Thus, we merged and integrated our myonuclei data with that of Sun et al., and generated a merged UMAP using our workflow (Figure 5B). This showed that the Unclassified clusters A and B myonuclei did not cluster in our merged UMAP but were spread in the *Myh1, Myh2* and *Myh4* clusters (Figure 5B-5C). This is consistent with the observation that myonuclei expressing *H19, Myh8, Myh3* and *Atf3* constitute a subset of body myonuclei (Figure S6). We confirmed that among specific biomarkers identified in the Sun et al. analysis some (*Runx1, GM28653, Dlg2*) are expressed by Sun et al. myonuclei and some (*Runx1, Gm28653*) by 7OV myonuclei (Figure 5B-5C) (Figure S6). We further observed on this merged UMAP an UM cluster belonging to the Sun et al. data set, that we designated as UM-Sun (Figure 5B). The UM-Sun myonuclei cluster mainly contains nuclei originating from Sun experiments, but also includes UM-Ctl and UM-7OV myonuclei, suggesting that all these myonuclei are transcriptionally similar, as exemplified by their expression of *Plcb4* and *GM28653* (Figure 5C). The expression of the biomarkers *Ebf1, Rian* and *Meg3* (Figure 5C) (Figure S7) is more specific of UM-Sun.

**Fig. 5.**
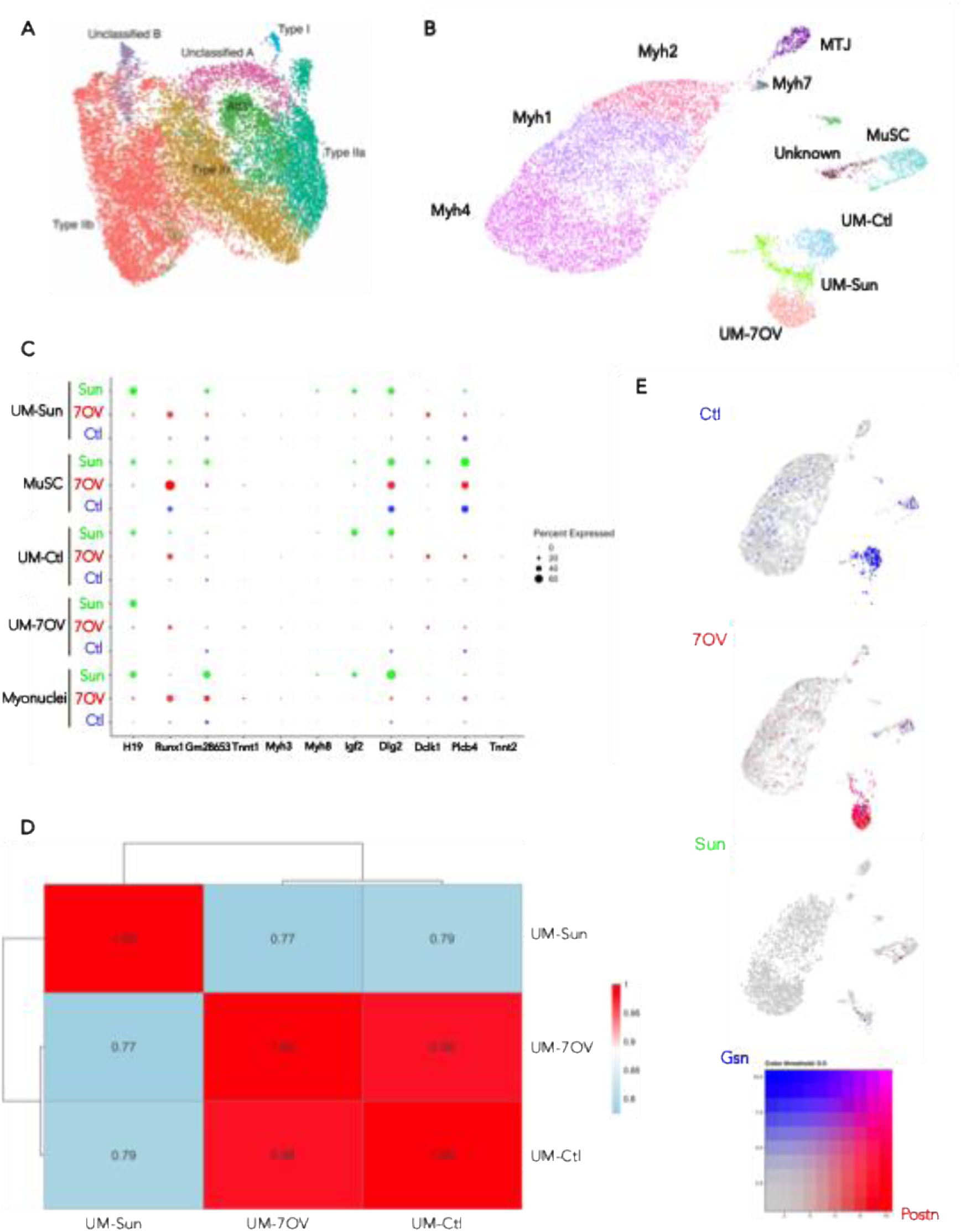
Unknown population of myonuclei is present in 7OV plantaris muscle. **A** Uniform Manifold Approximation and Projection (UMAP) visualization represents all myonuclei of Sun et al. **B** UMAP representing all the myonuclei of Sun et al. and of WT and mTmG mice. **C** Dotplot depicting biomarkers of Sun et al. in each myonucleus-type of Sun et al., WT and mTmG mice. **D** Transcriptional similarity between UM-Sun, UM-Ctl and UM-7OV cluster in Sun et al., WT and mTmG myonuclei. **E** Dimplot with expression of UM-7OV biomarker (*Postn*, periostin; red) and UM-Ctl biomarker (*Gsn*, gelsolin; blue) in all myonuclei from Sun et al., WT, and mTmG datasets. The UM-7OV cluster, marked by *Postn* is enriched specifically in the 7-day overloaded condition. The UM-Ctl cluster, marked by *Gsn* is present in control myonuclei. Grey nuclei express neither marker. The bicolor representation allows simultaneous visualization of the two distinct unknown myonuclear populations and highlights their condition-specificity: Gsn⁺ UM-Ctl in homeostatic muscle, and Postn⁺ UM-7OV emerging under mechanical overload.

This study also showed that *H19* is detected in all populations derived from the Sun experiments, and that *Runx1* marks all myonuclei populations from the 7OV experiments, suggesting that all nuclei may respond to a specific stress during the experiment, whathever on the method used to extract them.

We also performed an analysis of the transcriptional similarity between these three UM clusters (Figure 5D). The visible difference in correlation between UM-Ctl and UM-7OV (r = 0.90 in Figure 4, and r = 0.98 in Figure 5) is due to the use of a different UMAP (the one in Figure 4 did not include the results of Sun et al.) and therefore a clustering that may differ. With correlation values of 0.77 and 0.79, we further suggest that these clusters are transcriptionally related and likely originate from the same cell type, albeit with some differences. Finally, we demonstrated that the UM-Sun cluster had their own biomarkers *Ebf1*, *Rian* and *Meg3* and did not express *Postn* and *Gsn* like UM-7OV and UM-Ctl (Figure S7) (Figure 5E).

All of these results allowed us to conclude that UM-7OV are not newly fused myonuclei (Unclassified A and B in Sun et al.). We were also able to identify an UM cluster in the data from Sun et al. with transcriptional expression similar to our UM. This result supports the presence of UM myonuclei within the *plantaris*.

### Putative function of UM-7OV

To describe the hypertrophic process in more detail, we compared 7OV myonuclei with Ctl myonuclei (all myonuclei except UM-7OV and UM-Ctl, referred as body myonuclei). This comparison enabled us to identify 277 genes that were differentially expressed in body myonuclei under the two conditions. We then performed a Gene Ontology analysis on these differentially expressed genes to identify upstream regulators and canonical pathways predicted to be activated or inhibited. Twelve canonical pathways were predicted to be up-regulated and 8 down-regulated, and 35 upstream regulators were predicted to be up-regulated and 69 down-regulated. Many of the pathways involved in the remodelling of extracellular matrix elements are predicted to be activated in OV myonuclei (Figure 6A). In addition, MRTF2A, MRTF2B (SRF co-factors) and *YAP*, three key transcriptional transducers of mechanical signals, appeared among the upstream regulators predicted to be the most activated and have been implicated in the control of muscle mass (Figure 6B) [19,20,29,30]. The p53 protein, whose phosphorylation, stability and nuclear accumulation are induced by intense exercise was also predicted to be activated by OV [31].

**Fig. 6.**
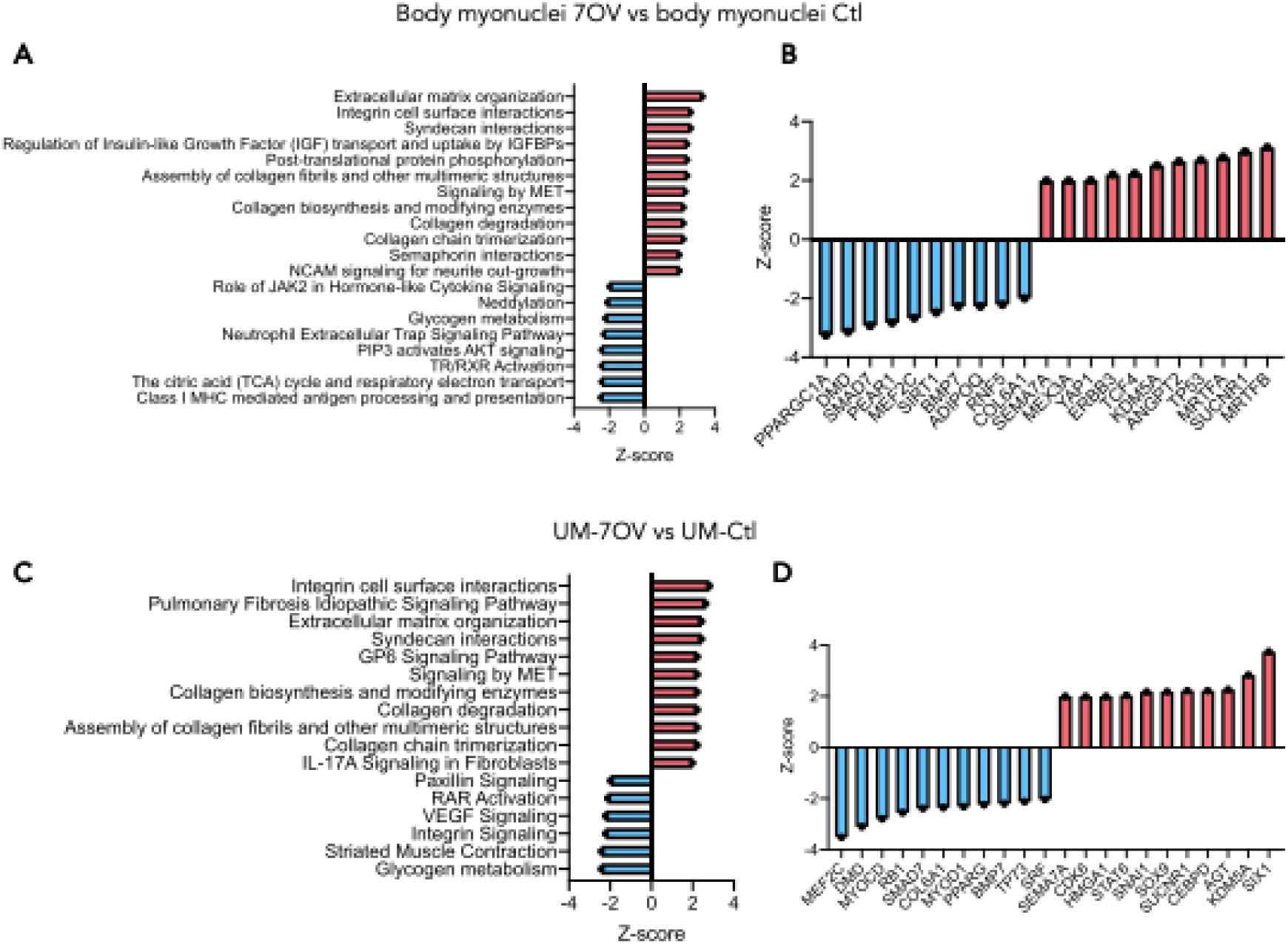
7OV myonuclei have a specific gene signature. **A** Predicted inhibition or activation of canonical pathways in 7OV body myonuclei compared to Ctl body myonuclei. **B** Predicted inhibition or activation of upstream regulators in 7OV body myonuclei compared to Ctl body myonuclei. **C** Predicted inhibition or activation of canonical pathways in UM-7OV compared to UM-Ctl. **D** Predicted inhibition or activation of upstream regulators in UM-7OV compared to UM-Ctl.

Trajectory analysis revealed a connection between the UM-Ctl and UM-7OV clusters, suggesting their proximity and transcriptional similarity. We also aimed to identify the signaling pathways involved in these myonuclei during hypertrophy by comparing the genes expressed in UM-7OV and UM-Ctl. A total of 136 genes were differentially expressed between UM-7OV and UM-Ctl, which is less than the number of genes found after comparing body OV and Ctl myonuclei (277 genes). After performing GO analysis, we obtained 11 predicted up-regulated and 6 down-regulated canonical pathways, and 23 up-regulated and 48 down-regulated upstream regulators. Again, many of the predicted activated canonical pathways were involved in extracellular matrix remodeling, including collagens (Figure 6C). Surprisingly, muscle contraction signalling was predicted to be inhibited in UM-7OV, whereas some upstream regulators predicted to be activated in UM-7OV are involved in inhibiting regeneration, differentiation and adult myogenesis, such as HGMA1, STAT6 and SNAI1 [32–34], while MYOD1 was predicted to be inhibited (Figure 7D). Additionally, in contrast to what was found for body myonuclei during OV, no upstream regulators involved in mechano-transduction were predicted to be activated. Furthermore, SRF, whose transcriptional activity responds to mechanical cues, was predicted to be inhibited (Figure 6D). We did not find any upstream regulators involved in hypertrophy or in the signaling pathways known to be involved in this process, with the exception of SUCN1R1, which is involved in the change in fiber type during hypertrophy [35]. Taken together, these predictions suggest that UM and body myonuclei may respond differently to overload-induced hypertrophy. In addition, SIX1, a key transcriptional regulator in muscle development and myofiber diversity, was among the most strongly predicted upstream regulators to be activated. The activation of SIX1 and its cofactor EYA3 during plantaris muscle overload has previously been characterized [36], as has the induction of *Myogenin* expression, a downstream target of the SIX transcriptional complexes [37].

**Fig. 7.**
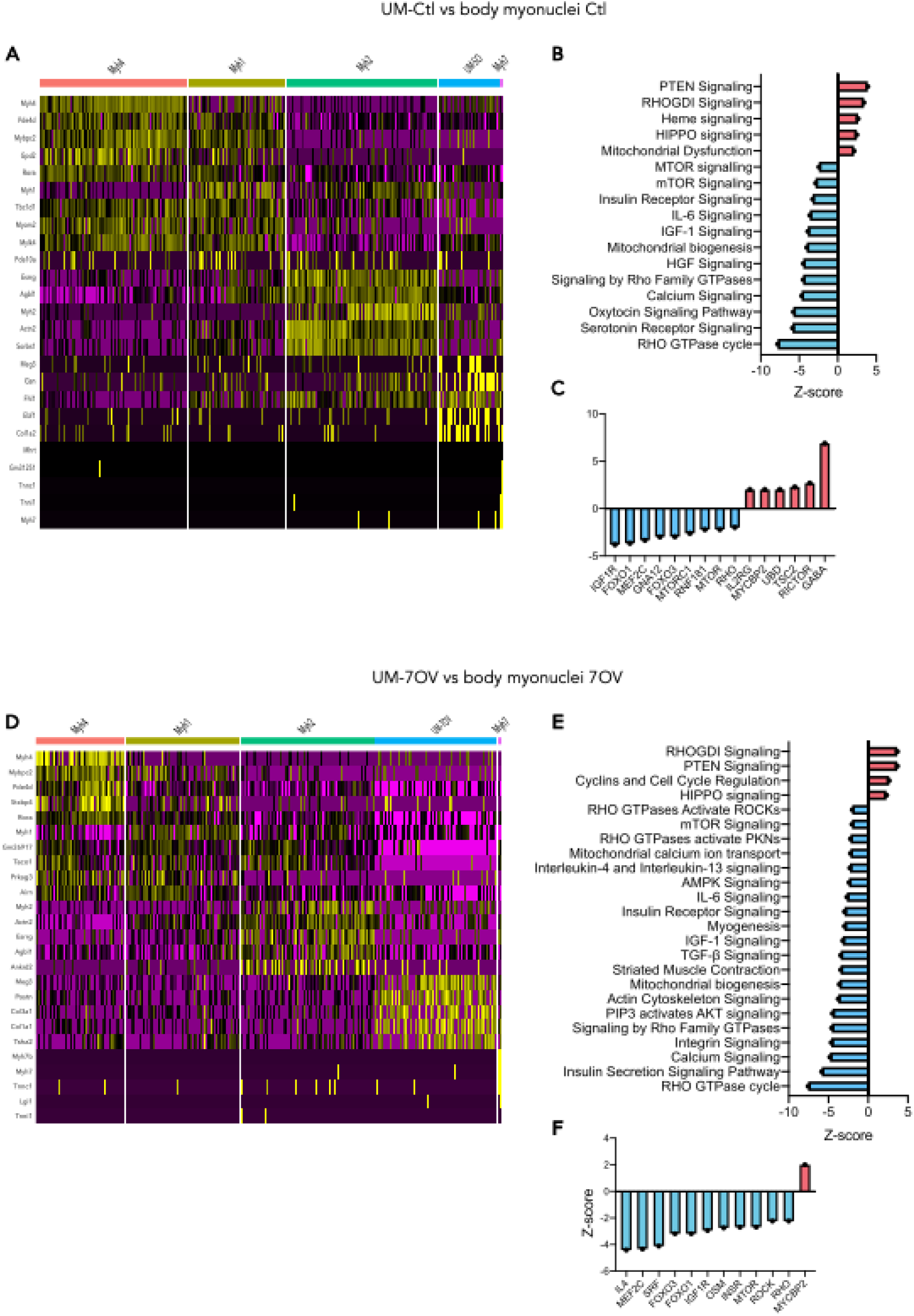
UM-Ctl and UM-OV have specific transcriptional expression. **A** Heatmap with top 5 biomarkers for *Myh1*, *Myh2*, *Myh4*, *Myh7* and UM-Ctl clusters. **B** Predicted inhibition or activation of canonical pathways in UM-Ctl compared to Ctl body myonuclei. **C** Predicted inhibition or activation of upstream regulators in UM-Ctl compared to Ctl body myonuclei. **D** Heatmap with top 5 biomarkers for *Myh1*, *Myh2*, *Myh4*, *Myh7* and UM-7OV clusters. **E** Predicted inhibition or activation of canonical pathways in UM-7OV compared to 7OV body myonuclei. **F** Predicted inhibition or activation of upstream regulators in UM-7OV compared to 7OV body myonuclei.

Finally, to further unravel the putative functions of UM-Ctl and UM-7OV, we compared their gene expression with that of the rest of the myonuclei (body myonuclei) under each condition, Ctl and 7OV. First, we compared UM-7OV with body OV myonuclei and UM-Ctl with body Ctl myonuclei. We found that 1348 genes were differentially expressed between UM-7OV and 7OV myonuclei, and 1567 between UM-Ctl and Ctl myonuclei (Figure 7A-7B). Using GO analysis of the differentially expressed genes, we identified the upstream regulators predicted to be specifically activated or inhibited in UM-Ctl and UM-7OV. For the comparison between UM-Ctl vs. Ctl myonuclei, we obtained 5 predicted up-regulated and 326 down-regulated canonical pathways, and 157 predicted up-regulated and 170 down-regulated upstream regulators. Signaling pathways such as mTOR and RhoA are predicted to be downregulated in UM-Ctl compared to other myonuclei in the basal state (Figure 7B). We can see that many upstream regulators of these two pathways are also predicted to be inhibited, allowing consistency between canonical pathways and upstream regulators (Figure 7C). However, RICTOR, an obligate component and activator of mTORC2, is predicted to be activated. This finding is difficult to reconcile given the simultaneous prediction of mTORC1 inhibition. However, mTORC1 and mTORC2 are distinct complexes with partially independent regulation and functions: mTORC1 primarily promotes protein synthesis and cell growth in response to nutrient and energy status, while mTORC2 mainly regulates the organization of cytoskeleton and cell survival via RICTOR and is predominantly activated by growth factors. The observed activation of RICTOR, and thus potentially mTORC2, and the inhibition of mTORC1 suggest separate upstream signals controlling these pathways in UM-Ctl compared to other myonuclei [38]. The upstream regulator with the highest Z-score was GABA. GABA is a factor that inhibits muscle function, is a muscle relaxant and appears to be up-regulated in ageing muscles. The predicted up-regulation of GABA in UM-Ctl suggest that UM-Ctl may dampen muscle contraction and act as a muscle relaxant. This aligns with findings that increased GABAergic activity can attenuate muscle contractile strength [39,40].

When comparing UM-7OV and 7OV body myonuclei, 4 canonical pathways were predicted to be up-regulated and 289 down-regulated, and 134 upstream regulators were predicted to be up-regulated and 172 down-regulated (Figure 7D). The results for both comparisons are quite surprising, as several key canonical pathways involved in hypertrophy are predicted to be inhibited, such as the mTOR, IGF-1, RhoA and actin cytoskeleton signaling pathways in UM-7OV (Figure 7E). The striated muscle contraction canonical pathway was also predicted to be down-regulated, as well as mitochondrial biogenesis and calcium signaling (Figure 7E). We then turned to the analysis of upstream regulators in the comparison of UM-7OV vs. body 7OV myonuclei. mTOR, IGF1-1 receptor (IGF1R) and insulin receptor (ISNR) activities were predicted to be inhibited, and a significant up-regulation of these pathways inhibitors such as MYCBP2 (e3-ubiquitin ligase) was predicted in UM-7OV (Figure 7F). However, other negative downstream effects of the mTOR pathway such as FOXO were predicted to be down-regulated, which is inconsistent with a putative inhibition of mTOR/Akt signaling (Figure 7F). This suggests that protein turnover in those myonuclei may be reduced, potentially leading to a slowdown in energy metabolism in the regions occupied by UM. Finally, we observed a significant down-regulation of the RHO signaling pathway and RhoA effectors such as SRF, MEF2C, IL4 and ROCK in UM-7OV compared to body 7OV myonuclei (Figure 7F).

## Discussion

A muscle fiber is a post-mitotic syncytium composed of myonuclei, which are accreted sequentially during development and myofiber growth, and reside in specific locations along/within the myofiber. Myonuclei are classified according to their genetic expression program and their location, such as MTJ myonuclei located next to the tendons and NMJ myonuclei located underneath motoneuron synapses [26,41]. How different myonuclei subpopulations may contribute to adult muscle adaptations such as overload-induced hypertrophy has been overlooked. Indeed, skeletal muscle hypertrophy has been studied considering that muscle tissue is a unique and indivisible entity, for example in terms of the signaling pathways involved. Moreover, with increased workload, a switch in gene expression towards a more oxidative metabolism accompanied by changes in genes coding for sarcomeric proteins such as *Myh1* and *Myh2* has been shown to occur at the fiber level, rather than at single-nucleus resolution [42].

In our study we identified myonuclei in the *plantaris* muscle during muscle homeostasis (UM-Ctl) and during the hypertrophic process (UM-7OV) that have a specific transcriptional program that distinguishes them from the rest of the myonuclei. These myonuclei express skeletal muscle markers, such as *Ttn* or *Myosins*, but also specific biomarkers such as *Gsn* for UM-Ctl and *Postn* for UM-7OV before and after OV, respectively. Importantly, trajectory studies suggested that UM-Ctl, became UM-7OV and UM-21OV with increased workload, and Gene Ontology analysis revealed that UM exhibited a distinct transcriptional response during hypertrophy compared to the other myonuclei.

Furthermore, UM-Ctl myonuclei do not correspond to the « unknown myonuclei » that we identified previously [9] nor to the new MTJ nuclei described by Kim and colleagues under resting conditions [10]. We initially assumed that UM-Ctl would be specific to the *plantaris* muscle, given that the majority of snRNA-seq studies have been performed on the *Tibialis anterior* (TA) or all hindlimb muscles. However, we did not identify the UM-Ctl cluster in the data from Sun and colleagues, where *plantaris* muscles were also analyzed. Moreover, when we analyzed the Sun data mixed to our data using our workflow, we observed a new population of myonuclei that we called UM-Sun. Those nuclei do not cluster in Sun data but clusterized close to our UM due to the existence of UM-7OV and UM-Ctl clusters in our data. Their transcriptional similarity permits the UM-Sun to clusterize. Therefore, the UM-Sun cluster was present and located close to UM-7OV, expressed *Meg3* like UM-7OV but did not express as *Postn* and have a correlation score r = 0.77 with UM-7OV and 0.79 with UM-Ctl, demonstrating their close transcriptional proximity [28]. These discrepancies observed could be due to differences in the nuclei purification/isolation protocols used, with Sun’s study performing specific sorting of myonuclei prior to snRNA-seq experiments. Another difference between these two studies is the invasiveness of the surgical technique used, with *Gastronemius* ablation in Sun et al. and tenotomy in our study [28]. Finally, there are differences in our workflow (among which doublet analysis and soupX) that can explain the absence of UM in Sun et al. data. Importantly, we were able to rule out the hypothesis of contamination or artefacts in our data sets and analysis because we performed two independent snRNA-seq experiments on OV and Ctl *plantaris* muscles from two different mouse strains (mTmG and WT), using both female and male mice, and obtained similar results and similar myonuclei clustering. Furthermore, our RNAscope data clearly identify *Postn* as a gene co-expressed with other specific muscle genes within myonuclei.

We next wondered whether the UM-7OV cluster might contain newly fused myonuclei resulting from the fusion of MuSCs with the growing myofibers. However, several lines of evidence argue against this hypothesis. First, the biomarkers of newly fused myonuclei identified by Sun et al. were not expressed in UM-7OV [28] and Figure 5). The evolution of the transcriptional programs of UM-Ctl, UM-7OV and UM-21OV revealed a close proximity between these clusters, suggesting the temporal transcriptional adaptation of the same myonuclei before (UM-Ctl) and after (UM-7OV and UM-21OV) overload-induced hypertrophy. This suggests that UM-Ctl in the basal state becomes UM-7OV during hypertrophy in response to increased mechanical signals and tension, and then reaches at an overloaded intermediate state UM-21OV, between UM-7OV and UM-Ctl, when muscle growth reaches a plateau [43].

The comparison of the UM-Ctl and UM-7OV clusters revealed that only a small number of genes were differentially expressed between the two clusters. Nevertheless, this comparison predicted the upregulation of several signaling pathways associated with extracellular matrix remodeling in UM-7OV. ECM remodeling is essential for myofiber hypertrophy, and this process is regulated by coordinated signaling from myofibers, satellite cells, and macrophages, involving specific molecular mediators and cellular interactions that underpin these adaptations [44,45]. In addition, we have previously demonstrated the action of RhoA on the ECM *via* activation of Erk1/2 and consequently the ECM regulatory factors MMP9, MMP13 and ADAM8 [27]. ECM remodeling pathways were also predicted to be activated during OV. These findings are in line with several studies that have shown ECM rearrangement during OV and the need for proper ECM integrity to maintain myofiber mechanical properties and sustain growth [27,46–48]. Interestingly, one of the genes over-expressed and characteristic of UM-7OV is *Postn*. *Postn* encodes a secreted ECM protein that binds to integrins and that has been shown to be involved in inhibiting muscle regeneration [49] and myogenesis [50], suggesting a role in muscle adaptation processes. The presence of *Postn* and its up-regulation suggest that its expression in the UM-OV may be required to modulate the hypertrophic response, potentially by influencing ECM remodeling and cellular signaling pathways associated with muscle growth [49]. Furthermore, many collagens are expressed in UM-7OV cluster, such as *Col1a1*, *Col5a1* and *Col3a1*. By upregulating the expression of so many extracellular matrix proteins, UM-7OV may play a crucial role in matrix remodeling during hypertrophy. This remodeling could affect fiber maintenance, as previously suggested, activation of MuSCs or their fusion with the growing myofibers [27]. Given periostin’s critical involvement in regulating cardiac hypertrophy and remodeling [51], studying *Postn* conditional mutant mice in skeletal myofibers could provide important insights into the respective roles of UM-7OV and periostin during hypertrophic processes. Whether increased SIX1 transcriptional activity in specific UM-7OV cells during muscle overload contributes to local extracellular matrix (ECM) remodeling—similar to processes observed during fetal development [52]—remains an interesting possibility.

Interestingly, GO analysis predicted the activation of key upstream regulators of mechanotransduction such as RHOA, MRTF and SRF during OV in body myonuclei but not in UM-7OV (vs. UM-Ctl). These predictions are consistent with previous studies that have reported the involvement of MRTFA and SRF in the control of muscle mass in response to mechanical cues. During disuse atrophy, MRTFA translocates outside the myonuclei and is therefore unable to activate SRF-dependent transcription which is important for muscle growth [19,30]. Furthermore, local and global mechanical stretch applied to myoblasts have been shown to induce the nuclear accumulation of MRTFA [53]. The fact that these mechanosensitive regulators are not predicted to be activated in UM-7OV, and that SRF is even predicted to be inhibited, highlights that UM myonuclei respond differently to OV compared to the rest of the myonuclei and do not seem to be coordinated with the other myonuclei in the myofiber [9].

The comparison of the UM-Ctl with Ctl body myonuclei clusters and the UM-7OV with 7OV body myonuclei cluster highlights a high level of differential gene expression for both, suggesting that UM have a specific transcriptional program within the plantaris muscle. Examining canonical pathways and upstream regulators revealed that UM-7OV and 7OV body myonuclei respond differently to hypertrophy, with a predicted downregulation of the mTOR and RhoA signaling pathways. These results are rather surprising, as their localisation at the tips of the myofibers suggested that they might be sensitive to mechanical signals. The predicted downregulation of the mTOR signaling pathway involved in hypertrophy suggests that UM-7OV are not the main players in increasing protein synthesis in the myofiber. However, our results also show that the mTOR inhibitor FOXO is predicted to be down-regulated in UM-7OV leading to a decrease of both activators and inhibitors of protein synthesis. Further studies are needed to determine whether mTOR signaling is indeed downregulated specifically at the tips of myofibers in plantaris muscles subjected to 7 days of overload, and to assess whether the observed increase in protein synthesis is regionally localized within the myofiber in a manner that promotes an increase in myofiber thickness rather than length. UM-7OV may locally down-regulate the canonical hypertrophic signals protecting the distal structure of the myofiber and avoiding the over-activation of protein synthesis and over-enlargement of the myofibers near the extremities. Our results therefore suggest that the mTOR pathway may be activated non-homogeneously along the fiber and that it may be mainly activated in the center of the myofiber, to allow enlargement of the myofiber mainly in this region. Further studies are required to confirm the non-homogeneous presence of mTOR activation along the fibers and the preferential enlargement of myofibers in their middle regions.

## Conclusion

Our results reveal the heterogeneity of myonuclei within the skeletal muscle syncytium and identify a specific population of nuclei localized preferentially at the ends of myofibers. These nuclei exhibit distinct transcriptional profiles compared to other myonuclei. We identified two key genes that characterize these tip nuclei: *gelsolin*, which marks their basal state, and *periostin*, expressed during hypertrophy. Our findings suggest that these nuclei transition from the basal to the hypertrophic state, enabling them to respond to hypertrophic signals. Importantly, these tip myonuclei maintain a distinctive transcriptional signature that sets them apart from central nuclei both at rest and during hypertrophy.

## Abbreviations

Ctl: control
Gsn: Gelsolin
HSA: Human skeletal actin
IP: intraperitoneal
MTJ: myotendinous junction
mTmG: membrane Tomato membrane
GFP: Musc muscle stem cells
Mut: mutant
Myh: myosin heavy chain
NMJ: neuromuscular junction
OV: overload
Postn: Periostin
SnRNA-seq: single nucleus RNA sequencing
TMX: tamoxifen
Ttn: Titin
UM: unknown myonuclei
UMAP: uniform manifold approximation and projection

## Declaration

### Ethics approval

Animal experimentation adhered strictly to the institutional guidelines for care and use of laboratory animals as outlined in the European Convention STE 123 and the French National Charter on the Ethics of Animal Experimentation (n°44083-2022121514506347, n°2024022718312758 and n°201809041512432, agreement n°C75-14-02). All animal procedures received approval from the French Ethical Committee of Animal Experiments CEEA - 034 and were performed in Institut Cochin animal core facility (Agreement A751402).

## Consent for publication

All authors have read and approved the final manuscript.

## Author Contributions

Designed experiments: Léa Delivry, Mathieu Dos Santos, Pascal Maire, Athanassia Sotiropoulos. Performed experiments: Léa Delivry, Stéphanie Backer, Maxime Di Gallo, Mathieu Dos Santos, Athanassia Sotiropoulos. Bioinformatic analysis: Léa Delivry, Alix Silver. Interpreted the data: Léa Delivry, Florian Britto, Pascal Maire, Athanassia Sotiropoulos. Wrote the manuscript: Léa Delivry, Pascal Maire, Athanassia Sotiropoulos. Pascal Maire and Athanassia Sotiropoulos supervised the project and acquired fundings.

## Acknowledgements

We thank D.Millay and C.Swoboda for meaningful discussions on bioinformatic analysis and for critical reading of the manuscript, Franck Letourneur, Sébastien Jacques and Brigitte Izac of the GENOM’IC facility, Thomas Guilbert and Pierre Bourdoncle of the IMAG’IC facility, Rémi Pierre and Marcio Do-Cruzeiro of the MOUST’IC core facility and Antoine Guéraud of the Institute Cochin for their technical assistance.

