## Supplementary Figures for "Identification of a new population of myonuclei during skeletal muscle hypertrophy"

Supplementary Materials include : 8 supplementary figures.

### Supplementary Figures

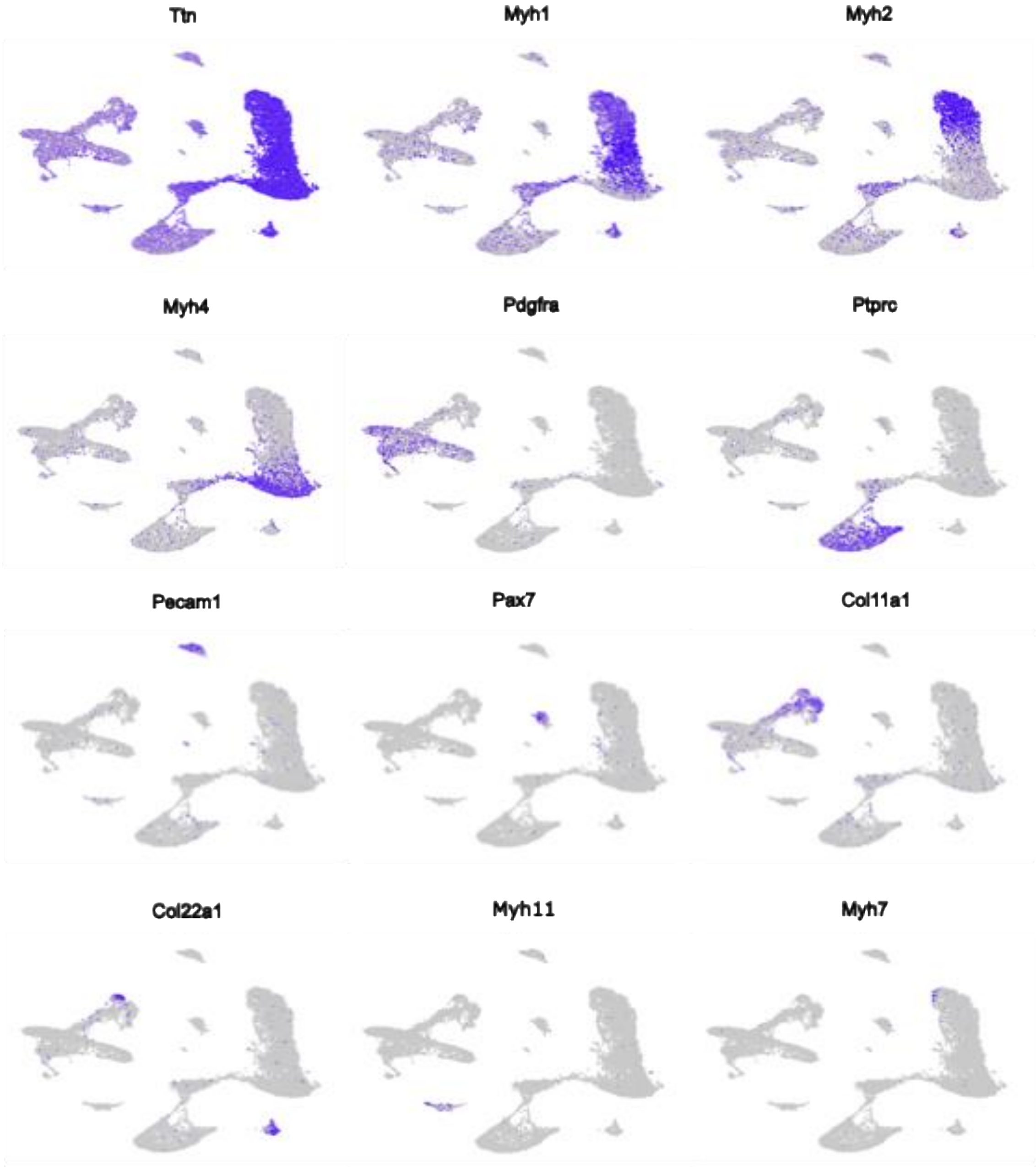


**Fig. S1** Gene expression in plantaris muscle from integrated snRNA-seq data.

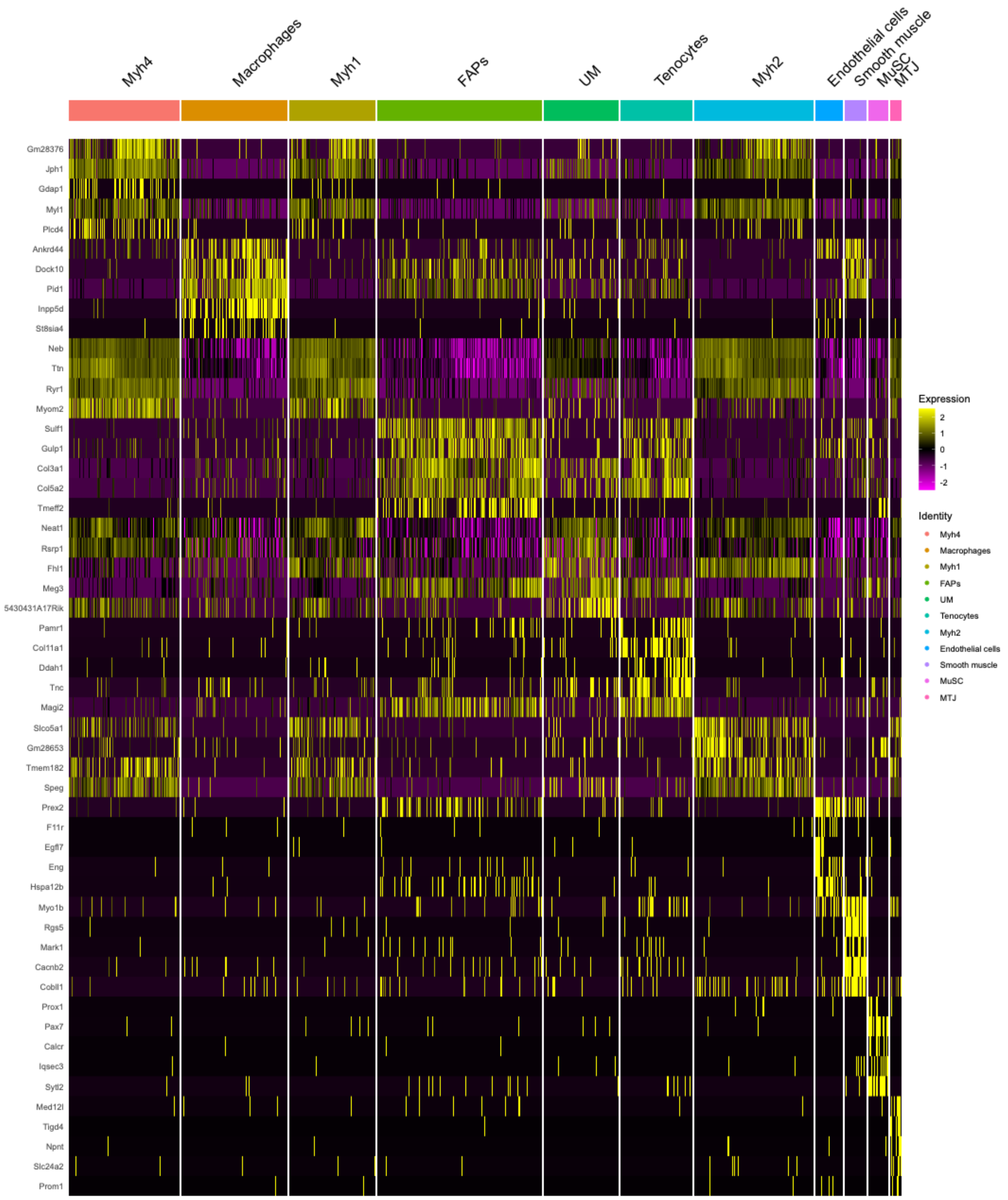


**Fig. S2** Heatmap of top biomarkers in each cluster of the Figure 1A UMAP.

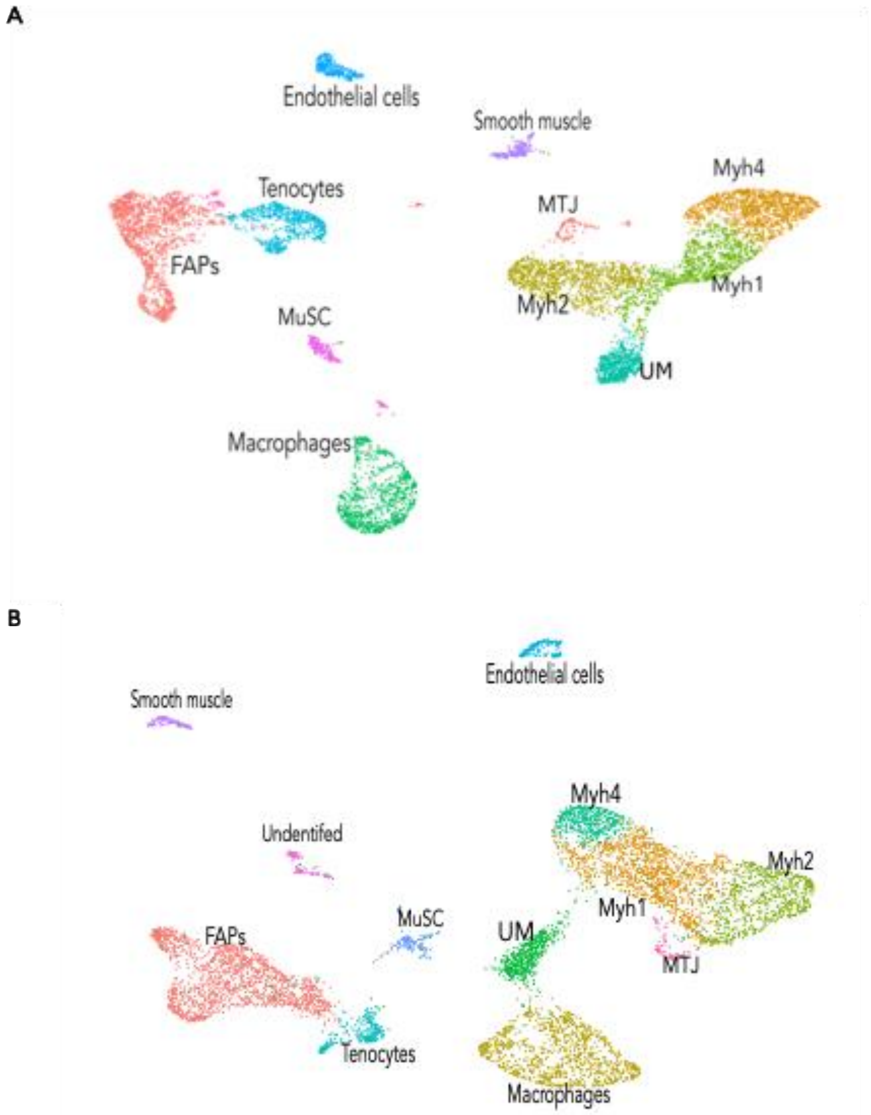


**Fig. S3** UMAP of each mouse model. **A** UMAP with only WT myonuclei SO and 7OV. **B** UMAP with only *ROSA*$^{mTmG}$*;Pax7*$^{CreERT2/+}$ (mTmG) SO and 7OV myonuclei.

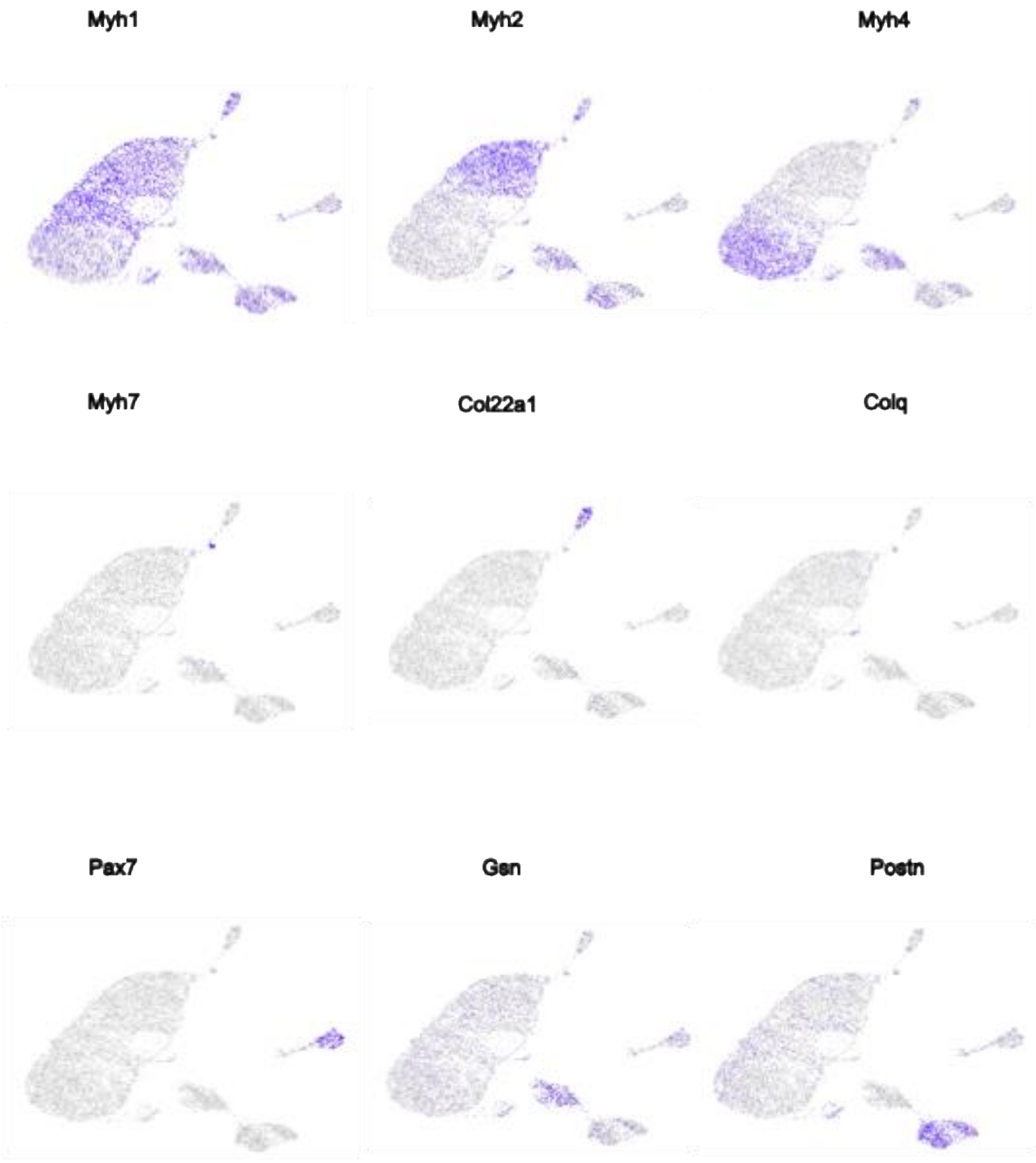


**Fig. S4** Gene expression of specific markers in myonuclei population from integrated snRNA-seq data.

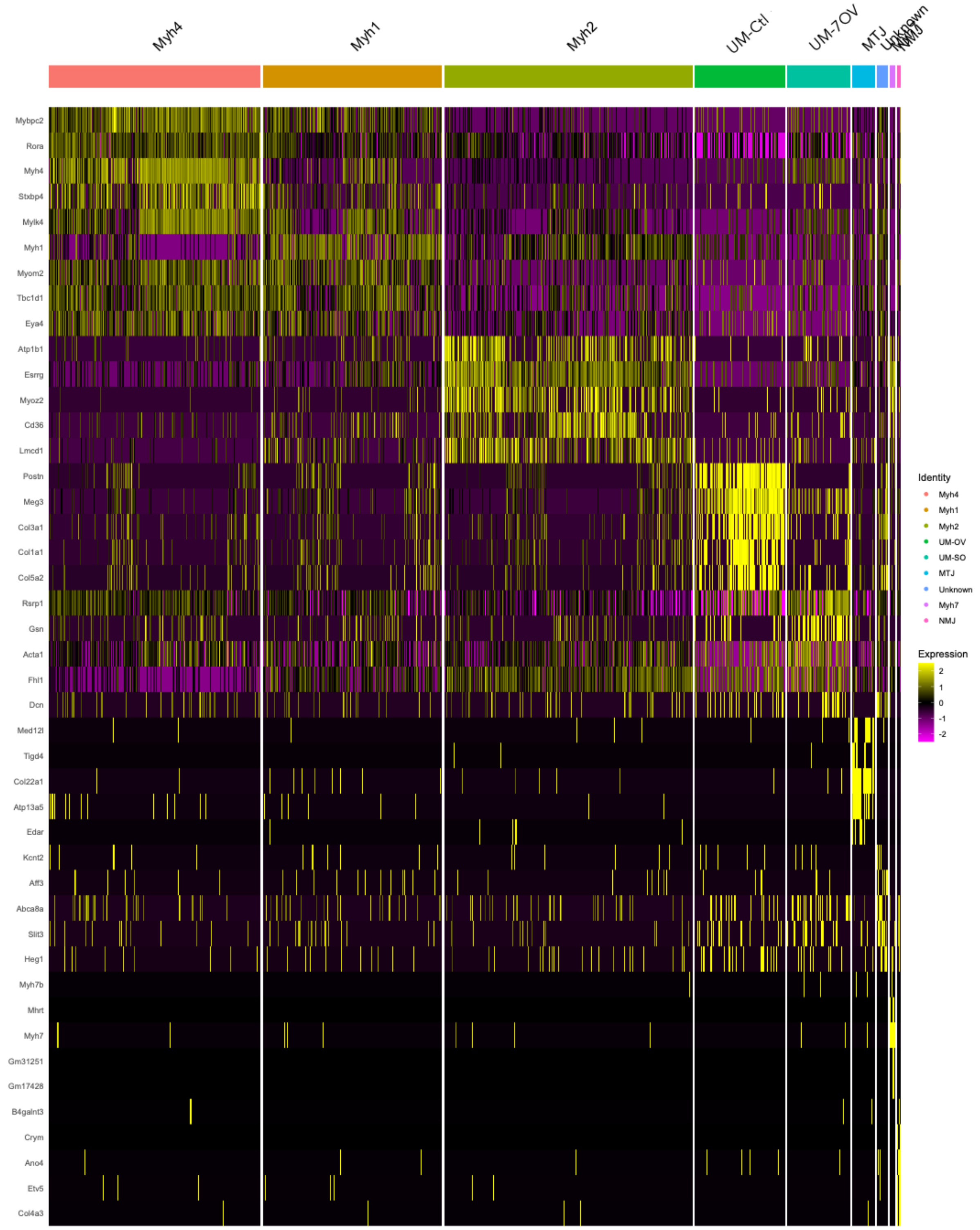


**Fig. S5** Heatmap of top biomarkers in each cluster of the UMAP of Figure 1B.

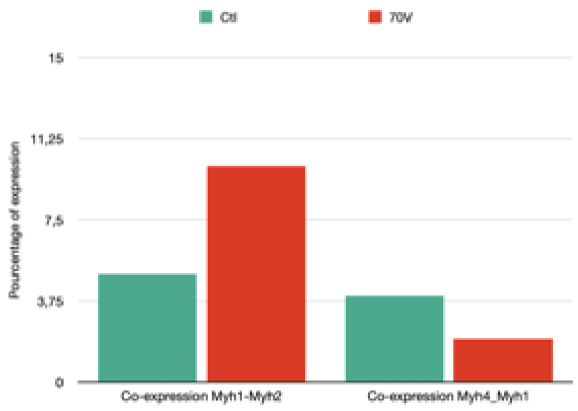


**Fig. S6** Co-expression of *Myosin heavy chain* genes in Ctl and 7OV myonuclei.

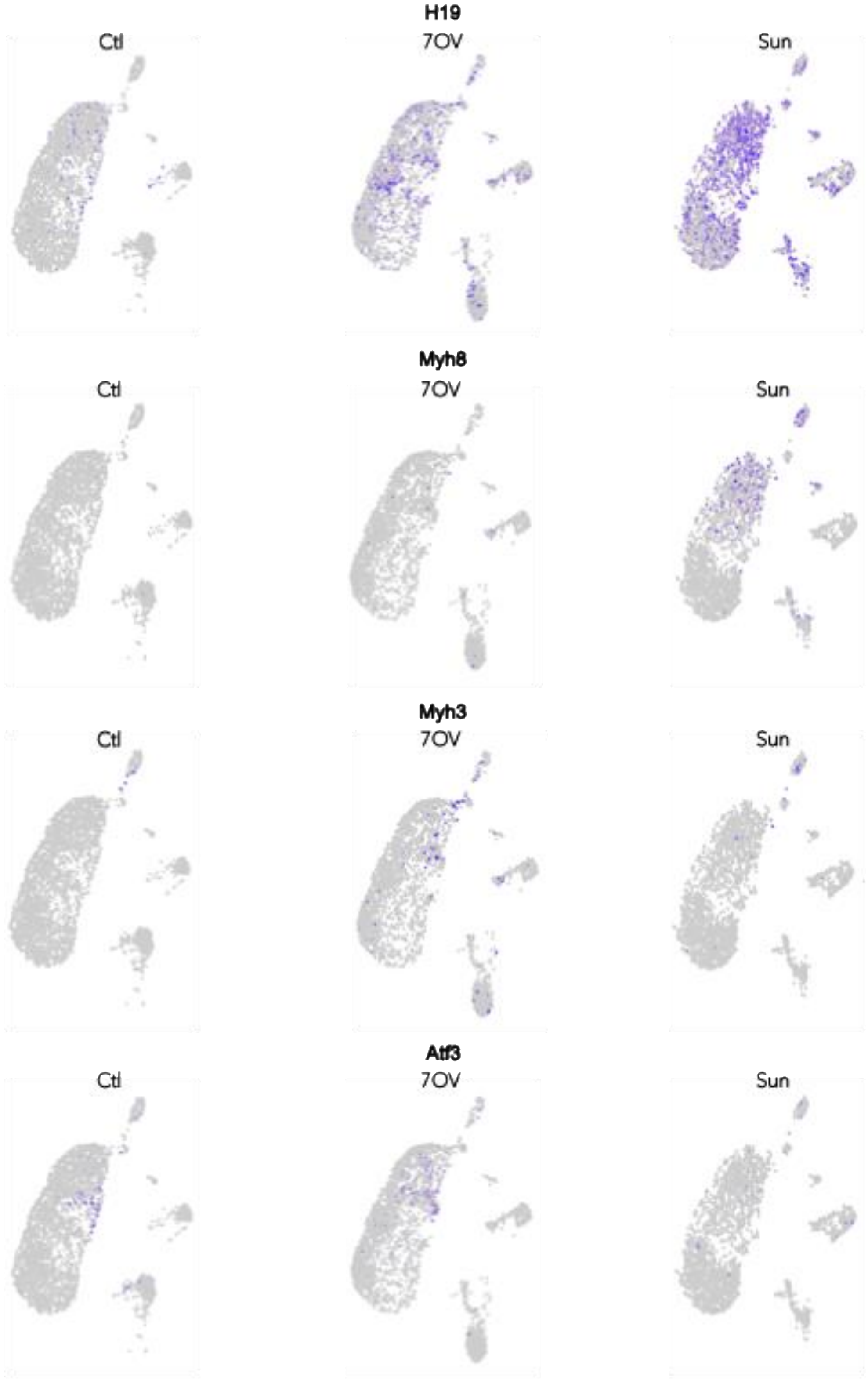


**Fig. S7** Gene expression of Sun et al. biomarkers in myonuclei population from merged integrated of our data and Sun et al. data.

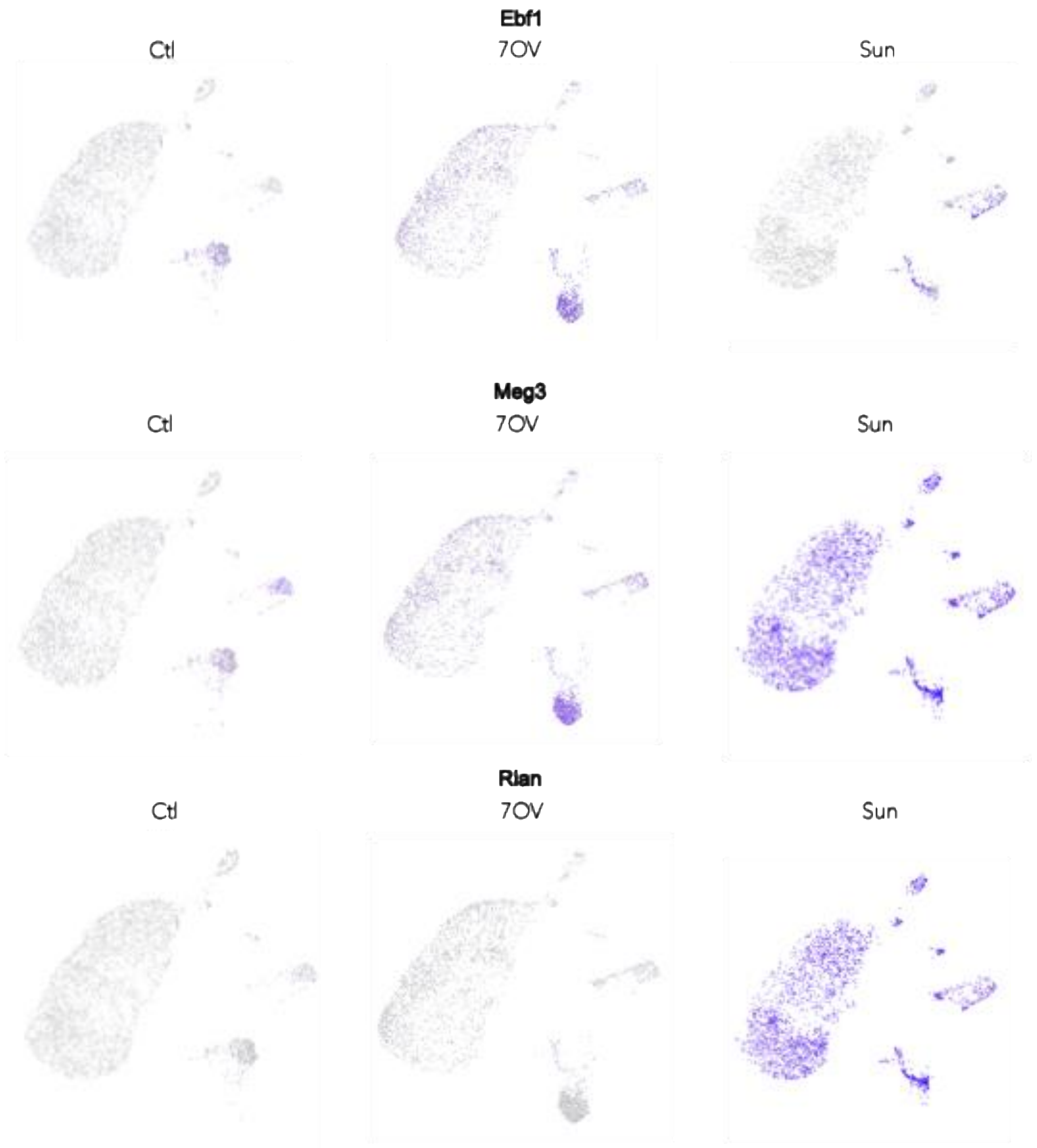


**Fig. S8** Gene expression of *Ebf1*, *Meg3* and *Rian* in myonuclei population from merged integrated of our data and Sun et al. data.
